# Ratiometric 4Pi single molecule localization with optimal resolution and color assignment

**DOI:** 10.1101/2021.10.30.466398

**Authors:** Jianwei Chen, Benxi Yao, Zhichao Yang, Wei Shi, Tingdan Luo, Peng Xi, Dayong Jin, Yiming Li

## Abstract

4Pi single molecule localization microscopy (4Pi-SMLM) with two opposing objectives achieves sub-10 nm isotropic 3D resolution with as few as 250 photons collected by each objective. Here, we developed a new ratiometric multi-color imaging strategy for 4Pi-SMLM which employed the intrinsic multi-phase interference intensity without increasing the complexity of the system and achieved both optimal 3D resolution and color separation. By partially linking the photon parameters between channels with interference difference of π during global fitting of the multi-channel 4Pi single molecule data, we showed on simulated data that the loss of the localization precision is minimal compared with the theoretical minimum uncertainty, the Cramer-Rao lower bound (CRLB).

---

In the past decade, single molecule localization microscopy[1,2] has revolutionized the field of biological imaging by improving the resolution of conventional fluorescence microscopy by more than an order of magnitude, achieving nanometer scale imaging resolution that is capable to resolve nanostructures of molecular machinery inside intact cells[3,4]. In particular, imaging schemes incorporating two objective lenses to coherently detect single molecule fluorescence in a 4Pi geometry has demonstrated to improve the axial resolution dramatically, even surpassing the lateral resolution[5]. The high axial sensitivity is due to the fact that the interference phase of the 4Pi-PSF directly couples with axial position, resulting in the fast change of the intensity of the 4Pi point spread function (PSF) along axial direction. By comparing the intensity of single molecules in the three[6] or four[7–9] interference phase channels, one can get a very sensitive readout of a single molecule’s axial position. The resulting *xyz* resolution is about 1.4×1.4×8 times better than that achievable using the single objective system. Therefore, by combining SMLM with 4Pi detection, a near-isotropic resolution down to ∼10 nm has been achieved[6–8,10] even with dim fluorescent proteins, enabling the application of 4Pi-SMLM in studies of nanoscale of protein localizations[11,12].

Multi-color SMLM is crucial to investigate the spatial relationship and interactions among different biomolecules. To further utilize its superior resolving power, different multi-color strategies have been applied in combination with SMLM. The most straightforward implementation is to sequentially image molecules labeled with spectrally different dyes[13]. Another option is to use the same fluorophore to sequentially label the molecules of interest and image them in different cycles (*e*.*g*., DNA-PAINT)[14]. However, the acquisition time for each detection channel adds up, therefore the imaging time is extended and it is more vulnerable to the sample drift.

Ratiometric multi-color imaging can distinguish the color of spectrally similar single molecules by using the relative intensity information between two spectral channels splitted by a dichroic mirror[15]. This approach has several advantages over conventional multi-color imaging using spectrally distinct dyes: 1) Multiple best ‘blinking’ dyes for SMLM are far-red dyes; 2) Chromatic aberration is neglectable which is important for ultra-high resolution multi-color imaging; 3) Only one laser is utilized to perform simultaneous multi-color imaging. However, conventional 4Pi single molecule data analysis workflow using photometry between different interference phase channels is not suitable to perform ratiometric color assignment as the intensity information is used to determine the interference phase. The first approach of using ratiometric multi-color imaging in 4Pi-SMLM is to employ the intensity difference between the P and S polarization channels by placing a dichroic mirror in front of the camera at a specialized angle[7]. Recently, Zhang *et al*. solved this problem by using an additional camera to collect the salvaged fluorescence reflected by the main excitation dichroic mirror, and determined the color by the intensity ratio between the salvaged fluorescence and normal fluorescence[16]. However, it comes with the cost of increased hardware complexity.

Latest developments in 4Pi-SMLM have introduced spline interpolated PSF models [17]to directly fit the 4Pi single molecule data instead of photometry-based methods, allowing an analytic model to accurately describe the fringe like 4Pi-PSF[18,19]. With either spline interpolated IAB-based or phase retrieved experimental 4Pi-PSF model, people have achieved CRLB in both simulated and experimental 4Pi single molecule data using direct model fit. Since the 4Pi single molecule imaging is intrinsically collected in multi-interference channels, we hypothesize that one can perform regular ratiometric multi-color imaging among different interference channels, thus achieving multi-color 4Pi-SMLM without adding additional detection channels.

**Fig. 1** is the optical design proposed for our ratiometric mutli-color 4Pi-SMLM. Similar to the regular 4Pi microscopy, the emitted fluorescence photons are collected by both objectives and interferes with themselves at a 50/50 beam splitter (BS). The transmitted and reflected interference channels separated by the BS has the opposite interference with a phase shift of π. The localization precision at the positions close to the intensity peak and valley is poor as the intensity hardly changes. To introduce more interference channels other than only two interference channels with a phase difference of π, 3 or 4 interference channels are normally used in the detection of 4Pi-SMLM. Here, we followed the work by Huang *et al*.[8] and introduce 4 interference channels by separating the p and s polarization channels with a phase shift of π/2. In order to perform ratiometric multi-color imaging between these channels, we then insert filters in one or two of the interference channels after BS to create an intensity difference among channels for spectrally different dyes as shown in **Fig. 1**. We finally use the intensity difference of each molecule to determine its color information.

**Fig. 1.**
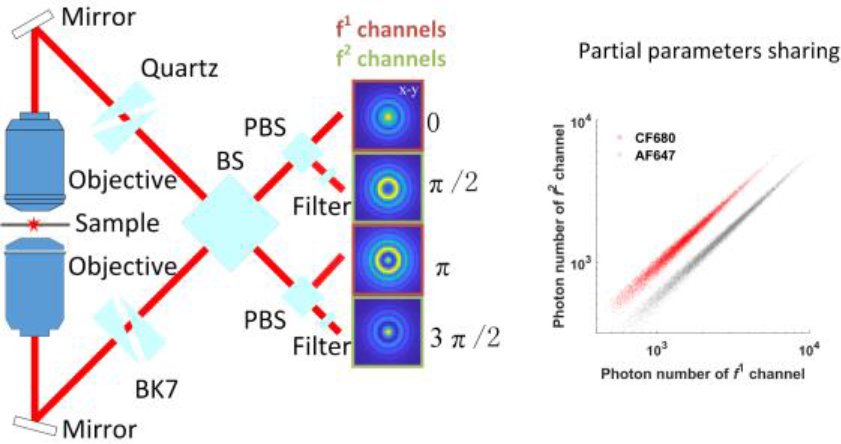
Schematic of the ratiometric multi-color 4Pi-SMLM employing the intrinsic multi-phase interference channels detection. The emitted fluoresce is collected by upper and lower objectives. Before self-interfering at the beam splitter (BS), the phase of p and s polarized fluorescence are shifted by two modified Babinet–Soleil compensators so that 4-phase inference channels (0, π /2, π and 3/2 π) are detected simultaneously. The 4-phase channels are then separated by a polarizing beam splitter (PBS) and collected at different regions of one camera. One or more filters are inserted in different interference channels to create intensity difference for spectrally different dyes.

To achieve the theoretical limit of the 3D localization precision (CRLB) for 4Pi-SMLM, we adapted the experimental IAB-based 4Pi-PSF model[18]. The experimental 4Pi-PSF model is defined as *P*(*x, y, z, φ*) = *I* + *Acos*(*φ*) + *Bsin*(*φ*). Here, *I*(*x, y, z*), *A*(*x, y, z*) and *B*(*x, y, z*) are slowly varying functions of *x, y, z* and independent of interference phase *φ*. We then divide the interference channels into two classes: 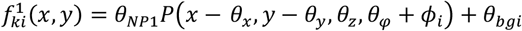 and 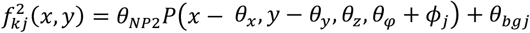. Here, 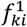 and 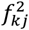 are the expected intensity value of *k*th pixel at position (*x, y*) in *i*th and *j*th channel respectively. The photonnumbers in *f*^1^ channels are with the same value *θ*_*NP*1_, while the photon numbers in *f*^2^ channels are with the same value *θ*_*NP*2_. In this work, we insert a filter in the *f*^2^ channels to create intensity difference between *f*^1^ and *f*^2^ channels so that ratiometric multi-color imaging can be performed. *θ*_*x*_, *θ*_*y*_, *θ*_*z*_ and *θ*_*ϕ*_ are the *x, y, z* positions and interference phase of the emitter, respectively. They are global parameters and the same for all channels. *θ*_*bgi*_ and *θ*_*bgj*_ are the constant background photons per pixel over the extent of the PSF in *i*th and *j*th channel, respectively. *ϕ*_*i*_ and *ϕ*_*j*_ are the corresponding relative phase shift in each channel which is controlled by the Babinet–Soleil compensators. The estimation of the parameters *θ* is achieved by using maximum likelihood estimation (MLE) based nonlinear optimization whose cost function across different channels is defined as: 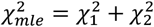. Here, 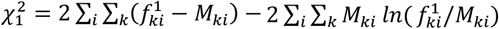 and 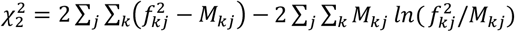. 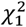 and 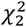 are the negative log likelihood function for *f*^1^ and *f*^2^ channels respectively. By minimizing 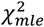, we obtain the maximum likelihood for the Poisson process.

We then evaluate the color separation of our proposed multi-color 4Pi-SMLM approach using simulated data. For comparison, we chose the same dyes as Zhang *et al*. for the three-color super-resolution imaging (CF680, DL650, DY634)[16]. The experimental single molecule photon distributions of these dyes are shown in **Fig. 2(a)**. To simulate the 4Pi single molecule data, we employed a full vectorial PSF model[20] and coherently added up the counterpropagating electrical fields from the upper and lower objectives. Here, an ideal 4Pi-PSF was simulated for both the upper and lower objectives with an NA of 1.35. The refractive indices of immersion medium and sample medium are both 1.40. The refractive index of the cover glass is set to be 1.518. The emission wavelength is 668 nm. An additional 60 mλ astigmatism was added to both upper and lower PSF. The same parameters are used throughout this work unless noted otherwise. We then decomposed the 4Pi-PSF into IAB-based 4Pi-PSF model which was used to fit the 4Pi single molecule data subsequently. 10,000 molecules were simulated with *x* and *y* positions randomly distributed within −1 to 1 pixels around the center of fitting window and *z* positions randomly distributed within −600 nm to 600 nm around the focus.

**Fig. 2.**
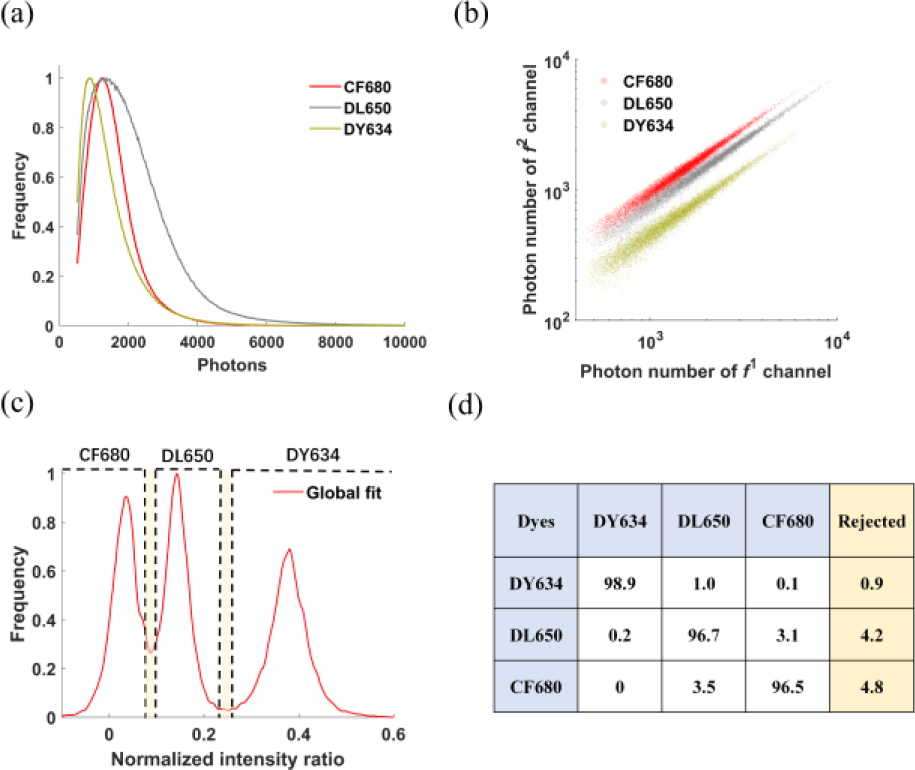
Color separation of ratiometric 4Pi-SMLM by partially linking the photons among different interference channels during fitting of multi-channel 4Pi single molecule data.(a) Experimental photon distributions of CF680, DL650 and DY634. (b) Scatter plot of the returned photons in *f*^1^ (0 and π) versus *f*^2^ (π/2 and 3/2 π) channels by IAB-based 4Pi-PSF model fit. (c) Normalized intensity ratio between *f*^1^ and *f*^2^ channels by partially linking photons in *f*^1^ and *f*^2^ channels separately. The molecules were assigned to 3 different colors based on the intensity ratio threshold by the 2 boxed region (molecules in the boxed regions were rejected. Left to right: CF680, DL650 and DY634,). **d**, The cross-talk (in %) of the 3 dyes using our proposed ratiometric multi-color 4Pi-SMLM method.

We then employed our global fit algorithm[21] to simultaneously fit 4 channels with *x, y, z* and phase as global parameters. The fitted photons of *f*^1^ versus *f*^2^ channels are shown in **Fig. 2(b)**. We then calculated the normalized intensity ratio *r* between *f*^1^ and *f*^2^ channels: *r* = (*θ*_*NP*1_− *θ*_*NP*2_)/((*θ*_*NP*1_+ *θ*_*NP*2_)). The histogram of r is shown in **Fig. 2(c)**. We assigned the molecules to 3 different colors based on the r threshold indicated by the 2 boxed regions. Molecules within the boxed regions are rejected. As shown in **Fig. 2(d)**, the crosstalk (in %) of the 3 dyes is less than 4% while the rejection ratio of the molecules is below 5%.

In the 4Pi-SMLM imaging, the intensities between different phase channels are normally highly correlated. Therefore, the first approaches to localize the *z* positions of 4Pi single molecules were to employ the intensity ratio between different channels to determine the interference phase[6–8]. We hypothesized that unlinking the photons between channels could lose information content and thus reduce the localization precision. To investigate the influence of our approach to the localization precision, we calculated the 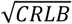 of the *x, y, z* positions within *z* range between 600 nm below and above the focus. Here, we compared the localization precision with photons linked/unlinked in all channels and partially linked between channels. Both 3 and 4 interference phase channels were investigated. As expected, when the photon parameter is fitted individually for each channel, the 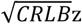 is much worse compared to that when the photon parameter is fitted globally across different channels (**Fig. 3 (a)** and (**b)**, yellow and red lines). The 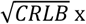 and 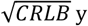 were not affected by different photon linking schemes (**Supplement Fig. 1**).

**Fig. 3.**
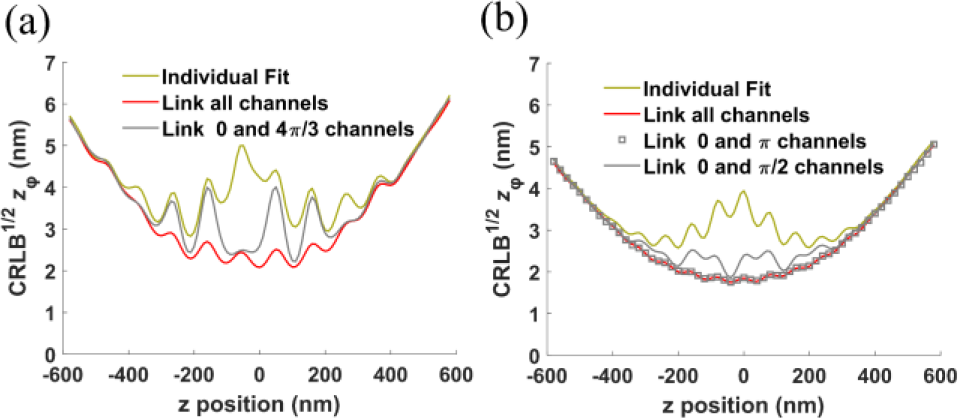
Comparison of the 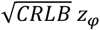 by partially linking of photons between different multi-phase interference channels in 3 (**a**) and 4 (**b**) phase interference 4Pi-SMLM. For each phase channel, 1,000 photons/localization and 20 background photons/pixel were used.

We then further investigate whether the CRLB could be improved by partially linking the photons within *f*^1^ and *f*^2^ channels. During fit, the photons in *f*^1^channels are shared as the same parameter *θ*_*NP*1_ while the photons in *f*^2^ channels are shared as the same parameter *θ*_*NP*2_. For the 3 interference phase channels 4Pi-SMLM imaging (0, 2π/3, 4π/3), we insert a filter in one of the channels (*i*.*e*.,4π/3). Therefore, 0 and 2π/3 channels are grouped as *f*^1^ channels and 4π/3 channel is classified as *f*^2^ channel. We then compared to the 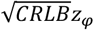 under this partially linking scheme to the fully linked condition. Although the 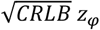 was improved compared to that when photon is individually fit, there are some peak positions where the 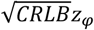 is much worse compared to that if the photons are fully linked between channels (gray line in **Fig. 3(a)**). For the 4 interference phase channels 4Pi-SMLM imaging (0, π/2, π, 3π/2), we partially linked the photons between channels in two different ways. In the first method, 0 and π/2 channels were grouped as *f*^1^ channels, π and 3π/2 channels were grouped as *f*^2^ channels. Similar to the 3 interreferences phase case, there are some peak positions of 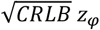 (gray line in **Fig. 3(b)**). In the second method, 0 and πchannels were grouped as *f*^1^ channels, π/2 and 3π/2 channels were grouped as *f*^2^ channels, Surprisingly, we found that the 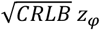 are almost the same compared to that when the photonsare fully linked (squaresin **Fig. 3(b)**). We therefore used this linking scheme for our ratiometric 4Pi-SMLM imaging strategy.

We further systematically evaluate the 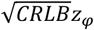 for differentphase shift Δϕ relative to π/2 between the p and s channels to find an optimal Δϕthat could achieve the theoretical best localization precision (**Fig. 4**). We compared the mean 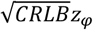 of *z* positions between ± 200 nm around the focus with 4 different parameter sharing schemes: link/unlink photons in all channels, partially link the photons in 0 and π channels, partially link the photons in 0 and π/2+Δϕ channels. We found that the mean 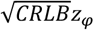 by partially linking the photons in 0 and π channels is the same as that by linking photons in all channels for all Δϕ (red line and squares in **Fig. 4**). When Δϕ = 0, mean 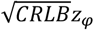 reached its minimal value. It indicates that the localization precision achieves optimal value when the phase shift between p and s channels is π/2.

**Fig. 4.**
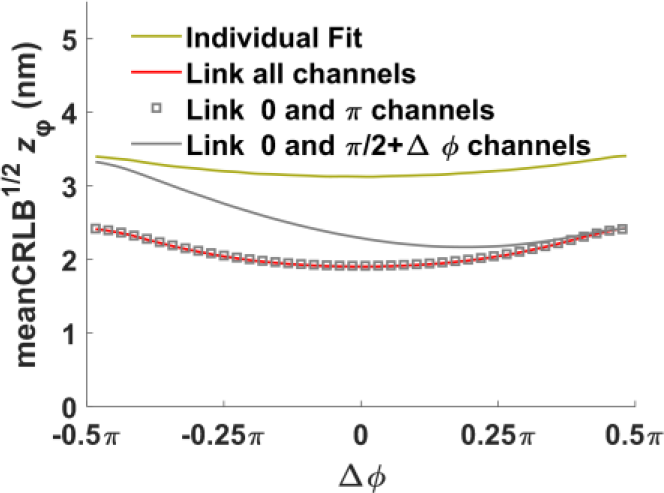
Theoretical minimum uncertainty of *z*_*φ*_ as a function of the phase shift between p and s polarization channels in the 4 interference phases 4Pi-SMLM imaging.

Since the ‘salvaged fluorescence’ method with 4Pi-SMLM system has been successfully applied for two and three fluorophores imaging simultaneously, we compared it to our proposed ratiometric 4Pi-SMLM method. Instead of using the salvaged fluorescence reflected by the main dichroic (ZT405/488/561/647rpcv2, Chroma) to distinguish the color, we choose a dichroic to maximize the detected photons as shown by the dashed line in **Supplement Fig. 2** (ZT405/488/561/640rpcv2, Chroma). The transmittance using new dichroic compared to the original dichroic can be found in **Supplement Table1**. The transmission fluorescence for the new dichroic was much higher compared to that for the original dichroic for DY634, AF647 and DL650 separately.

In order to create an intensity difference between *f*^1^ and *f*^2^ channels, we additionally insert a bandpass filter (ET710/75x, Chroma) to *f*^2^ channels. After inserting the filters, the photon ratios between *f*^2^ and *f*^1^ channels (*θ*_*NP*2_ /*θ*_*NP*1_) for CF680, DL650 and DY634 are 0.94, 0.75 and 0.45 respectively. For the ‘salvaged fluorescence’ method, we linked all the parameters in all four channels to get an optimal localization precision. For our new ratiometric 4Pi-SMLM method, we partially linked the photons in channels with a phase shift of π. As shown in **Fig. 5**, due to higher photon collection efficiency and optimized parameter sharing scheme, our proposed ratiometric multi-color 4Pi-SMLM approach outperformed the ‘salvaged fluorescence’ method in all four commonly used dyes for ratiometric multi-color SMLM imaging.

**Fig. 5.**
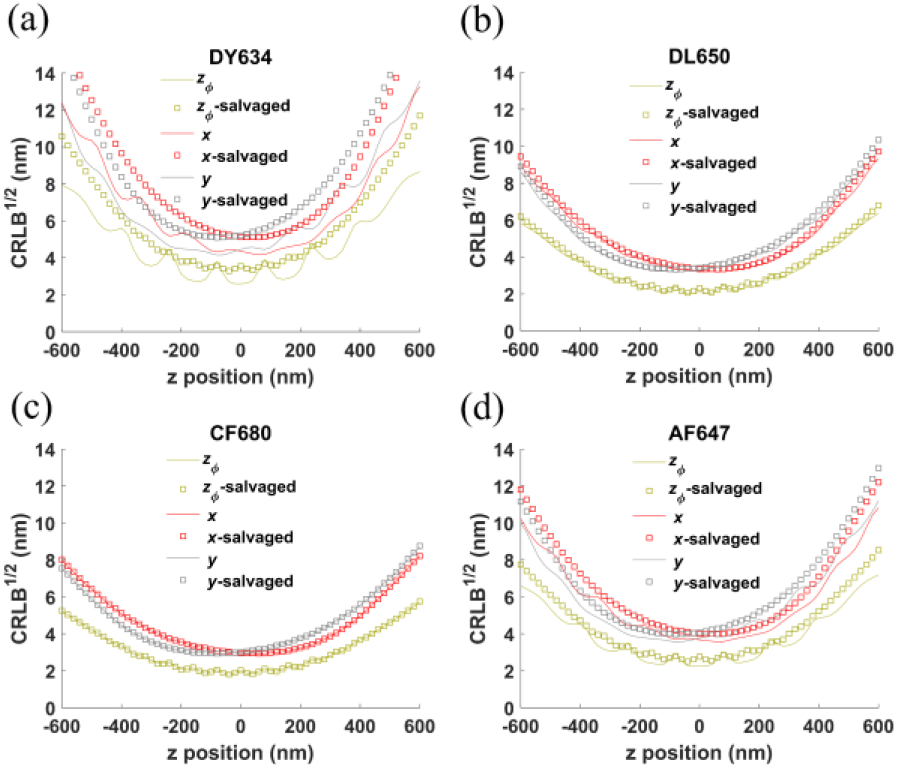
Comparison of the 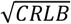 using partially linked ratiometric and salvaged fluorescence multi-color imaging for different dyes: (a) DY634; (b) DL650; (c) CF680; (d)AF647. For each phase channel, 1,000 photons/localization and 20 background photons/pixel were used.

In conclusion, we found the photon number of different interference channels is highly correlated in 4Pi-SMLM and strongly affects the localization precision. Our new ratiometric multi-color 4Pi-SMLM method could achieve good color assignment efficiency with minimal loss of information content within the multi-channel 4Pi-PSF, and therefore achieving optimal multi-color localization precision for 4Pi-SMLM. Especially for the four channel 4Pi-SMLM imaging, we found that the optimal resolution is obtained when the phase shift between p and s polarization channel is π/2. By partially linking the photons in channels with phase shift of π, almost no information content was lost as indicated by the 3D CRLB compared to the fitting scheme that links all parameters across all channels. Compared to the multi-color 4Pi-SMLM using ‘salvaged fluorescence’, our proposed methods could achieve higher photon collection efficiency and thus better localization precision without adding hardware complexity. Moreover, the partially linking global fitting algorithm used in this work was implemented in GPU, which is more than 38 times faster than the CPU code (**Supplement Fig. 3**). We believe that this work will make multi-color 4Pi-SMLM much easier to be implemented in the existing 4Pi microscopy, and thus make better use of its ultra-high 3D resolution imaging to investigate the spatial relationship between different biomolecules.

## Data Availability

The code used in this workis open source on GitHub (https://github.com/Li-Lab-SUSTech/Ratiometric-4Pi).

## Supplemental Document

See Supplement 1 for supporting content.

