## Supplement 1 for "Ratiometric 4Pi single molecule localization with optimal resolution and color assignment"

**Supplement Fig. 1**

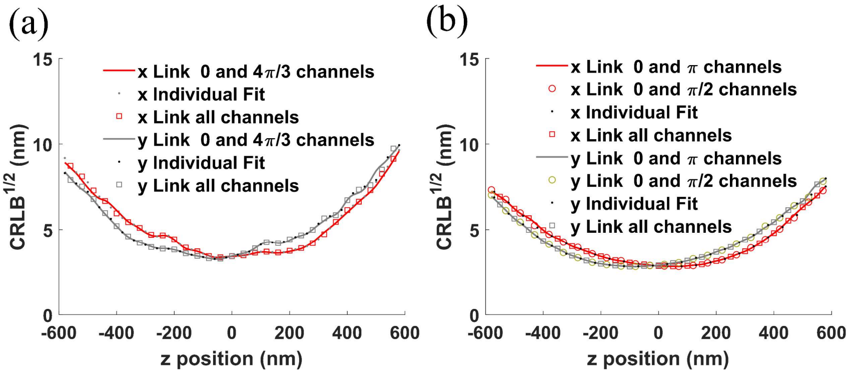


Comparison of the $\sqrt{CRLB}\ x$ and $\sqrt{CRLB}\ y$ by different parameters linking schemes for 3 (a) and 4 (b) phase interference 4Pi-SMLM. For each phase channel, 1,000 photons/localization and 20 background photons/pixel were used. We compared the localization precision with photons linked/unlinked in all channels and partially linked between channels. Both 3 and 4 interference phase channels were investigated. As shown in the **Supplement Fig. 1**, the $\sqrt{CRLB}$ x and $\sqrt{CRLB}$ y were not affected by different photon linking schemes.

**Supplement Fig. 2**

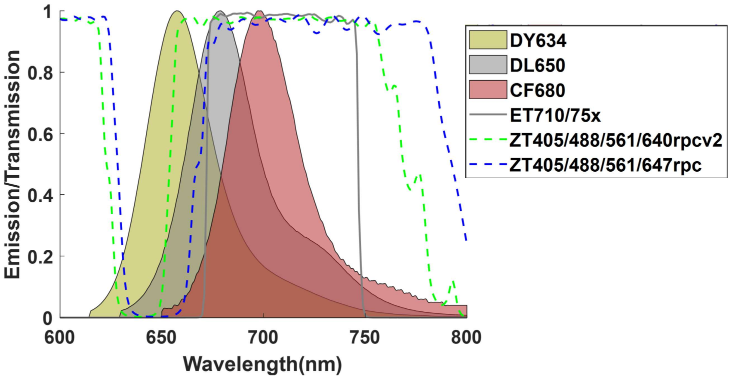


Emission spectra of three dyes (DY634, DL650 and CF680) and transmission spectra of the dichroic filter (ET710/75x, ZT405/488/561/640rpcv2 and ZT405/488/561/647rpc Chroma). In our proposed ratiometric mutli-color 4Pi-SMLM system, we use ET710/75X and ZT405/488/561/640rpcv2 to create photon difference in different channels. For the 'salvaged fluorescence' method, ZT405/488/561/647rpc was used. More fluorescence was reflected in the 'salvaged fluorescence' method as shown in **Supplement Table. 1**.

**Supplement Fig. 3**

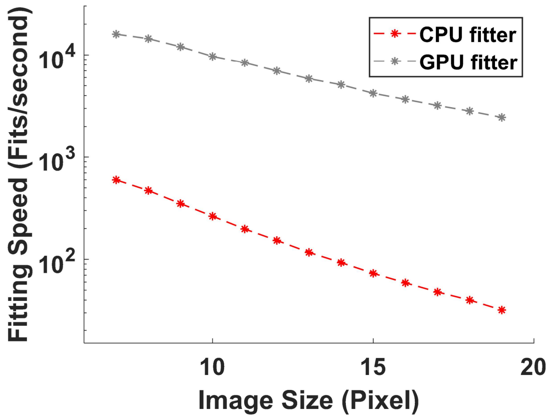


Computation speed of CPU and GPU global fitter for ratiometric 4Pi-SMLM as a function of image size. 4Pi image data with the same size was simulated for speed evaluation. In this case, the Intel Core i7-8700 @ 3.20 GHz and the NVIDIA GeForce RTX 3090 were used. The averaged fitting speed of GPU is about 38 times faster than that of CPU.

**Supplement Table. 1**

Transmittance of the dyes

| Dyes | Salvaged | Ratiometric |
|---|---|---|
| **DY634** | 43.2% | 53.8% |
| **DL650** | 77.9% | 81.4% |
| **CF680** | 93.6% | 93.9% |
| **AF647** | 60.2% | 68.4% |

Fluorescence photon transmittance for the 'salvaged fluorescence' method and our ratiometric mutli-color 4Pi-SMLM approach using the filters as indicated in **Supplement Fig.2**. The table shows that our ratiometric 4Pi-SMLM approach has higher photon collection efficiency compared to the 'salvaged fluorescence' method. Higher photon transmittance means that the system can collect more photons, thus achieving better single molecule localization accuracy.
